# A platform for the systematic interrogation of genetic interactions in human cells: 26 gene by genome-wide double knockout screens as a proof of concept

**DOI:** 10.1101/2025.08.04.665369

**Authors:** Andrew Chatr-aryamontri, Li Zhang, Ndeye Khady Thiombane, Vincent Archambault, Sébastien Carréno, Javier Di Noia, Ivan Topisirovic, Julie A. Lessard, Philippe P. Roux, Nicolas Pilon, Aaron Schimmer, Mike Tyers, Corey Nislow, Sylvie Mader, Brian T. Wilhelm, Thierry Bertomeu

## Abstract

Genetic interactions are typically studied by looking at the phenotype that results from disruption of pairs of genes, as well as from higher order combinations of perturbations. Systematically interrogating all pairwise combinations provides insights into how genes are organized into pathways and complexes to sustain cellular homeostasis and how interacting genes respond to stressors and external signals. Genetic interactions have been studied extensively in yeast, due, in part, to the availability of a systematic collection of gene knockouts, and the development of Synthetic Genetic Array (SGA) technology. In contrast, such approaches are more challenging in human cells and therefore comparable data for human cells is scarce. This study introduces an innovative approach to functionally characterize genetic interactions in human cells through CRISPR/Cas9 screens using a pooled genome-wide knockout library in NALM6 cells. By combining a single guide RNA (sgRNA) targeting the gene of interest (aka the query) in cells already infected with an inducible genome-wide sgRNA pool, it is possible to achieve near saturation of genome-wide double knockouts. We conducted 26 of these screens, which we term “gene by genome-wide” knockout screens. This approach can be rapidly performed, in part, because it bypasses the need to generate genotyped isogenic knockout clones. Data from these screens identified both expected and novel synthetic lethal and synthetic rescue interactions, demonstrating that this strategy is effective for large-scale genetic research in human cells. Additionally, we show that these GBGW screens can be combined with chemical perturbation to reveal new synthetic interactions that are not apparent without drug treatment. Finally, we show that cDNA overexpression can be incorporated with genome-wide knockouts to systematically explore gain-of-function scenarios. The complete dataset is accessible on the ChemoGenix website (URL: https://chemogenix.iric.ca).

## Introduction

A traditional approach to investigate the function of a gene is based on the *in vivo* inactivation or removal of the specific gene of interest followed by assays for its phenotypic impact. While this approach informs on the function of individual genes, it is subject to a high false negative rate, due in part to the fact that cellular functions are often accomplished by overlapping and partially redundant genes. The simultaneous inactivation of two genes may address this redundancy and, furthermore, can reveal the functional relationship between genes. When performed systematically, large-scale genetic interaction studies can yield comprehensive gene-gene interaction maps. In the yeast *S. cerevisiae,* the availability of complete arrayed collections of cells engineered with loss of function (LOF) of individual genes in both a and α yeast mating types (1) has enabled the systematic assessment of the impact of double gene knockouts on cellular fitness and the derivation of precise maps of gene-gene interactions controlling cellular metabolisms. The elaboration of equivalent genetic charts in human cells has been hampered by the lack of effective and scalable genome editing technologies and phenotypic screening platforms. The efficacy and specificity of the CRISPR/Cas9 knockout technology, combined with the throughput afforded by next generation sequencing (NGS) of cell populations transduced with genome-wide sgRNA pooled libraries, has successfully addressed this limitation (2,3). Integrating these individually robust methodologies has enabled the performance of comprehensive functional gene knockout screens in human cells, such as essentiality screens assessing the fitness deficits caused by the loss of gene function within the screened cell line (3).

In pooled CRISPR KO screens, individual sgRNA molecules are introduced into cells via lentiviral infection followed by integration into the host genome. The sgRNA directs Cas9 to the specific gene locus, resulting in its inactivation through DNA cleavage.

Quantifying the difference in sgRNA frequencies between the initial and the final cell pools indicates whether each gene knockout provides a selective, relative fitness advantage or disadvantage with respect to cell proliferation of the transduced pools. This format, however, is limited to the analysis of the impact of individual gene knockouts; adapting this technology and its downstream analysis to create a comprehensive genetic interaction network in human cells still poses considerable technical challenges. For example custom multiplexed sgRNA libraries can be effectively employed for investigating specific genetic interactions, particularly when focusing on a limited set of genes (4–14). However, the insights gained from this method are confined to the chosen genes, limiting the potential to uncover unforeseen relationships. This approach is not generalizable because of requirement to develop novel sgRNA libraries for each new genetic interaction space to be investigated.

Developing larger libraries that can cover broader interaction networks often comes at the expense of increased library complexity (number of guides per gene), which can introduce noise, ultimately diminishing the precision of the assay. To overcome the shortcomings of sub-pool library screens and to enable comprehensive genome interrogation, individual isogenic genotyped knockout clones can be generated and infected with a genome-wide CRISPR knockout lentiviral library (15). While this method guarantees an unbiased exploration of the genome represented in the library, it is labor-intensive and time-consuming which constrains the number of screen that can be performed, limiting its scalability. In addition, clone selection can also introduce biases in epigenetic landscapes, gene expression patterns and essential pathways affecting interpretation of the screens.

An alternative approach to screening individual clones that circumvents the creation of knockout clones involves transducing the cells with individual sgRNAs that target the query genes prior to pooled genome-wide library knockout induction with Cas9. This strategy has proved effective in testing both double and even triple gene knockouts (16). Here, we present a streamlined and reproducible method, where gene-by-genome double knockouts are constructed by integrating a sgRNA targeting a specific gene of interest into cells already infected with a pooled inducible genome-wide sgRNA KO library. To reduce variability that may arise from incomplete or lack of KO or gene-specific failure of selected sgRNAs chosen to produce knockouts, we performed two parallel experiments using two different sgRNA sequences per gene. In total, 26 genes were examined using this GBGW genome-wide double knockout methodology. The resulting dataset is the most extensive of this nature, comprising nearly 500,000 gene pair deletions. This resource can be accessed publicly on the website of the ChemoGenix CRISPR/Cas9 chemogenomic screening facility of the Institute for Research in Immunology and Cancer (IRIC) at the University of Montreal (https://chemogenix.iric.ca/).

## Results

### Experimental design

The experiments were conducted with a doxycycline-inducible Cas9 clone developed from the human pre-B leukemic NALM6 cell line, into which the EKO (Extended Knockout) pooled sgRNA library was integrated by viral infection. The EKO library comprises over 278,000 sgRNAs (∼10 sgRNAs per gene) targeting more than 19,000 RefSeq genes (3). This lentiviral library was expanded and aliquots stored in liquid N2, ensuring that the distribution of GW sgRNA frequencies remained consistent across all experiments performed from a single infection “batch”. This experimental design is a modification of the protocol utilized in our standard chemogenomic screens (17). For each screening round the EKO-infected cell pool was cultured for five days to yield 5 billion cells, a quantity of cells sufficient for up to 40 experiments. No notable depletion of sgRNAs targeting essential genes was detected in the absence of Cas9 activity (Suppl. Fig. 1), indicating that this period of outgrowth does not affect the pool representation. Following this expansion, the pool was divided into different samples that were each individually transduced with a different query sgRNA (day 0, Fig. 1A). As a proxy, in order to avoid time consuming individual gene target KO validation, we used two different guides in independent experiments for each target gene. Furthermore, based on known essentiality profiles, the impact of each individual KO on cell fitness provides an independent assessment of the sgRNA efficacy. After 8 days of hygromycin selection and seven days of Cas9 induction with doxycycline (with a three-day overlap between the selection and induction phases to reduce overall screening time), the cells were propagated for an additional 8 days to allow any fitness differences of the double mutants to manifest as enrichment or depletion. Because NALM-6 cells double every 24 hours, this time frame allows enough time for fitness differences to become apparent.

**Figure 1:**
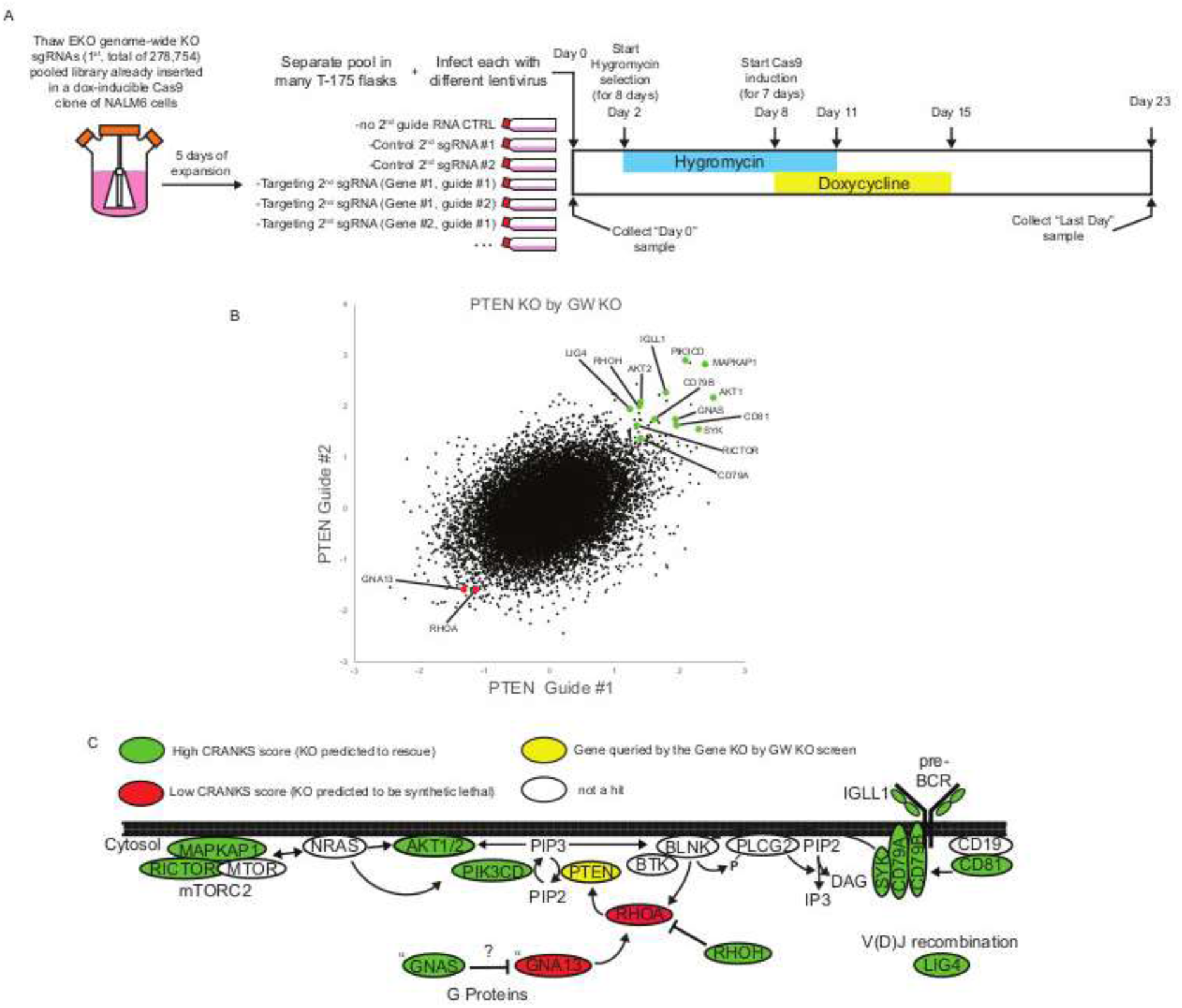
PTEN screen results. **A)** Timeline and overview of GBGW KO pooled screens. The uninduced GW EKO library is cultured over a period of five days. On Day 0 the cells are divided for infection with various lentiviruses to express a second sgRNA in all cells. Selection with hygromycin starts on Day 2 and continues for 8 days, overlapping with a 7-day induction of Cas9 using doxycycline. Following the selection and induction phases, the pools are expanded without doxycycline for an additional 8 days, culminating in collection on Day 23. **B)** The gene scores derived from the PTEN KO guide #1 and PTEN KO guide #2 screens are illustrated in the scatter plot. Select genes that exhibit a positive interaction with PTEN are indicated in green, while select genes that demonstrate a negative interaction are marked in red. **C)** Gene hits from screening PTEN KO effectively reflect the established biochemical interactions associated with PTEN. Additionally, they suggest a potential link in between PTEN and pre-BCR signaling pathways.

The EKO sgRNA sequences present in the genomic DNA extracted from cell pellets collected on day 0 and day 23 were amplified and sequenced by NGS to assess their abundance, determine frequency changes in target gene KO vs control populations and calculate gene depletion/enrichment scores as previously described (17).

### Gene by Genome Wide Screen Results

To evaluate the feasibility and performance of this approach, we targeted seven well-characterized genes (PTEN, TP53, MDM2, NPRL3, ADK, HMGCL, and PARP1), all of which are not critically essential in NALM6 cells (3). Average scores for each sgRNA in PTEN screens performed using two different levels of library depths (100 or 250 cells per sgRNA) were highly correlated (r=0.9) (Suppl. Fig. 2). Based on this observation, subsequent screens were performed on a broader selection of query genes based on research interests of partner labs using 100 cells/sgRNA. This provides a pragmatic balance that maximized multiplexing capabilities while minimizing costs. An additional 19 genes were screened, some of which have limited functional information available, including BANF1, CCNQ (FAM58A), CDADC1, CSE1L, DDX3X, DDX5, ERCC2, ERCC3, ETFDH, FAM172A (ARB2A), LONP1, MAPK14, PLCXD1, RBM26, SMARCD1, TOP2A, TOP2B, YME1L1, and ZNF451. The subsequent section highlights those screens that yielded noteworthy findings and which help demonstrate the potential of these GBGW screens. In each case we present only high level observations in the anticipation the readers will further mine the datasets.

**PTEN** is a lipid phosphatase that acts as tumor suppressor by attenuating PI3K/AKT signaling (18). The analysis of the PTEN screens with both sgRNAs demonstrates agreement among the top scores (Fig. 1B) and identified multiple genes that oppose PTEN’s function (including AKT1, AKT2, PIK3CD, and MAPKAP1) as rescue hits, i.e., their knockdown resulted in improved fitness (Fig. 1C). Interestingly, genes associated with pre-BCR signaling, such as SYK, CD79A, CD79B, CD81, and IGLL1 were found to act as rescues. These observations are consistent with the fact that NALM6 cells are a pre-B cell lineage and, as such, only express immature heavy-chain immunoglobulins associated with IGLL1, a pre-B cell receptor involved in proliferation and differentiation. Noteworthy synthetic lethal targets identified include LIG4, RHOA, and GNA13. LIG4 encodes a DNA ligase that plays a crucial role in V(D)J recombination, allowing maturation of NALM6 cells and the reconfiguration of signaling pathways that are downstream of pre-BCR (19). RHOA is known to activate PTEN, facilitating B cell proliferation and maturation (20). Additionally, GNA13 is an alpha subunit of a G protein that activates RHOA (21). Comparison of our PTEN results from NALM6 cells to those of Feng et al. (15) obtained from 293A cells revealed that both screens yielded a small number of hits, with a limited overlap between the two datasets (Fig. 2A). By way of example, our analysis did not detect SWI-SNF components ARID1A and SMARCB1, whereas Feng et al. failed to identify PIK3CD, AKT1, AKT2, and MAPKAP1 as enriched genes. Potential explanations for this limited conservation of hits can be attributed to the distinct biological characteristics of 293A vs NALM6 cells, as well as differences in the experimental design, including sgRNA libraries that vary in complexity and the use of isogenic vs pooled clones.

**Figure 2:**
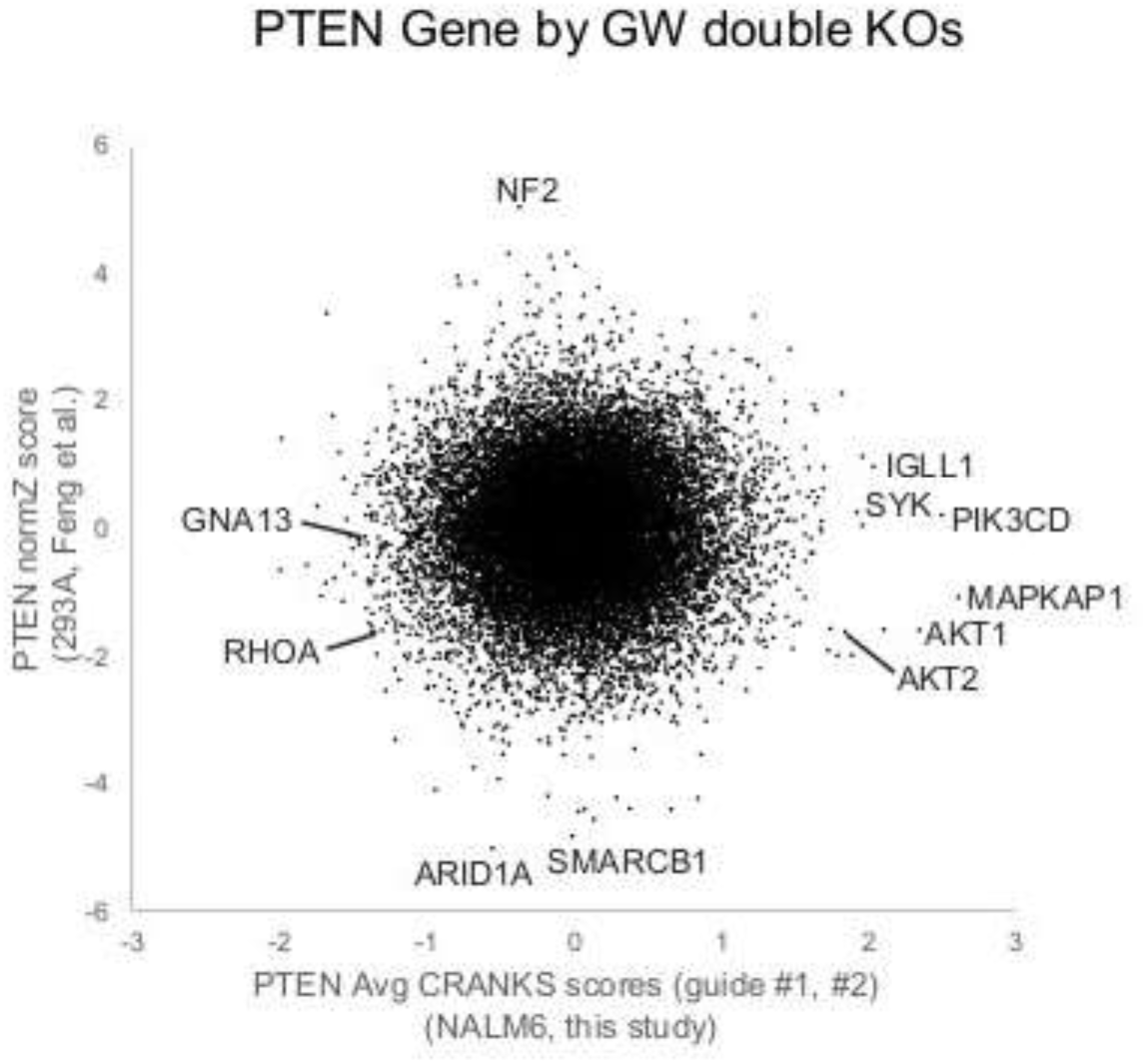
Comparative analysis of PTEN screens in NALM6 and 293A cells. Comparative analysis of our population level PTEN GBGW screens (average scores obtained with two distinct guides targeting PTEN) in NALM6 cells with the results from the screening of a PTEN KO clone derived from 293A cells described in Feng et al..

**MDM2** is an E3 Ubiquitin ligase that facilitates the ubiquitination and subsequent degradation of p53 (22). In NALM6 cells, MDM2 prevents the acute activation of p53 and is an essential gene (3). The top hit of the MDM2 screen was, as anticipated, the KO of TP53 which scored as a potent rescue hit (Fig. 3A). Another notable hit was unexpected; we found a strong genetic interaction of MDM2 with GPHN (Gephyrin), a microtubule-associated protein associated with synaptic biology, which acts as a scaffolding protein for glycinergic and certain GABAergic inhibitory synapses (23).

**Figure 3:**
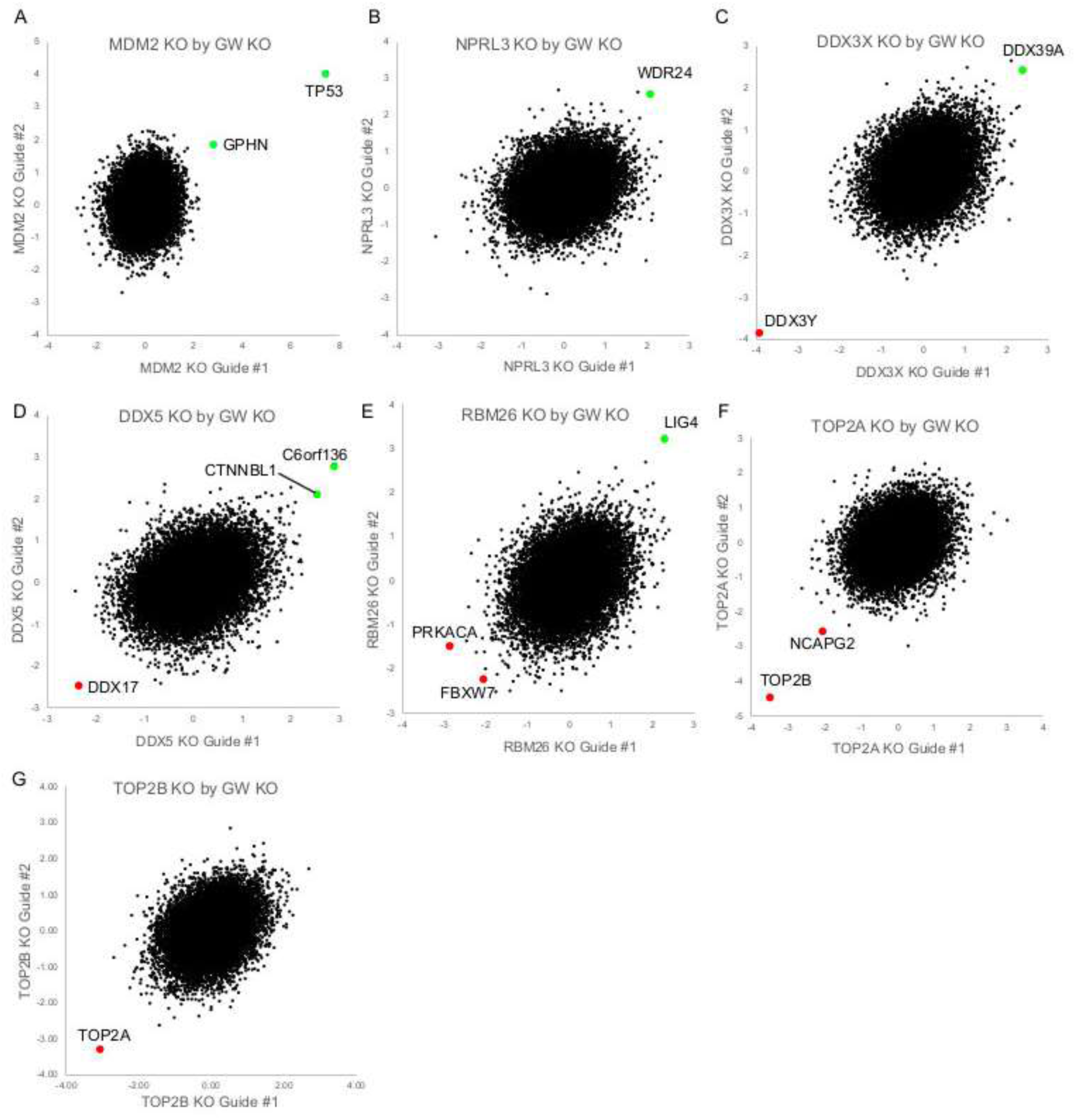
Gene by GW double KO screen results. Scatter plots display the CRANKS scores for all genes obtained by two distinct sgRNAs (guide #1 and guide #2) for each gene. Scores below zero indicate a depletion of sgRNA compared to the control, while positive scores reflect an enrichment of sgRNA. Selected hits are labelled and highlighted in green (rescues) or red (synthetic lethal).

**NPRL3** is a subunit of the GATOR1 complex, which inactivates the central metabolic regulator mTORC1 (24). This inactivation is reported to occur through enhancement of its GTP hydrolysis by the RRAGA/B heterodimer, rendering the heterodimer incapable of activating mTORC1 (25). The most significant rescue observed in the NPRL3 knockout screen was WDR24 (Fig. 3B), a subunit of the GATOR2 complex that activates mTORC1 by inhibiting the GTPase activity of GATOR1 (25). In NALM6 cells, WDR24 is classified as an essential gene, whereas NPRL3 is not (3). Because WDR24’s function is dependent on NPRL3, WDR24 may no longer be required for cell viability in the absence of NPRL3. No other GATOR2 subunits, such as MIOS and WDR59, emerged as significant hits, suggesting that they may act differently than WDR24.

**DDX3X** is an X-linked ATP-dependent RNA helicase that, among many other functions, promotes efficient translation of selected complex mRNAs (26). The most prominent synthetic lethal interaction identified in the DDX3X screen was with DDX3Y (Fig. 3C), its Y-linked paralog, which is known to be partially redundant with DDX3X (27).

Additionally, DDX39A, another dead-box RNA helicase involved in pre-mRNA splicing and nuclear export, emerged as the leading rescue in the DDX3X screen, suggesting that is has a potentially antagonistic biological role to DDX3X.

**DDX5** is a Dead Box RNA helicase that plays a crucial role in various biological processes, including regulation of splicing and transcription (28). The strongest synthetic lethal hit identified is DDX17 (Fig. 3D), a paralog of DDX5. This result is consistent with prior studies that suggest that DDX5 and DDX17 share partially overlapping functions (28,29). The rescue hit CTNNB1L is a part of the PRP19-CDC5L complex, which plays a crucial role in the activation of pre-mRNA splicing. This indicates a potential interaction between the two genes in the regulation of splicing processes. The specific role of the poorly annotated gene C6orf136 (which scored as a DDX5 interactor), in potentially counteracting the biological activities of DDX5 is less clear.

**RBM26** encodes a protein containing an RNA binding motif that, in association with Poly(A) Tail eXosome Targeting (PAXT), participates in the decay of nuclear mRNA (30) and in mRNA splicing via the recruitment of PABPN1 (31). The RBM26 gene was selected for screening because we previously identified it as a top hit in a chemogenomic screen with etoposide, a Topoisomerase 2 inhibitor (unpublished data). Although the specific function(s) of RBM26 in the DNA damage response pathway remain unclear, we reasoned that identifying its genetic interactors could be informative of its cellular roles. The most pronounced synthetic lethal hit was the cAMP-dependent Protein Kinase A gene (PKA). While one report indicates that PKA can phosphorylate XRCC6 (32), the relationship between PKA and DNA damage is not well understood. The second strongest synthetic lethal hit in the RBM26 knockout screen was the E3 ubiquitin ligase FBXW7, and the most significant rescue was the DNA ligase LIG4 (Fig. 3E). LIG4 is a component of the DNA-PK complex kinase, which includes PRKDC, XRCC4/5/6, DCLRE1C, and NHEJ1, and is involved in mediating Non-Homologous End-Joining (NHEJ) at DNA ends. FBXW7 is known to polyubiquitinate XRCC4, thereby enhancing its interaction with XRCC5 and XRCC6 (33). Additional work will be needed to determine if RBM26 influences the mRNA stability of DNA-PK components.

The topoisomerase 2 complex (Topo 2) resolves DNA entanglement by transiently inducing DNA double-strand breaks. In vertebrates, Topo 2 can function as **TOP2A** or **TOP2B** homodimers or as TOP2A/2B heterodimers. TOP2A is a core essential gene across multiple cell lines (https://depmap.org/), while TOP2B is non-essential. In the double genetic screens performed, TOP2A and TOP2B emerged as the most notable synthetic lethal interactions with each other (Fig. 3F-G), suggesting that TOP2B can partially compensate for TOP2A depletion and confirming that our approach can uncover functionally redundant bidirectional genetic interactions.

### Combining GBGW with Chemogenomics

Out of the 26 total GBGW screens, the above-described 8 yielded results that were straightforward to interpret, while the remaining 18 did not have particularly strong genetic interactors. This result was not unexpected given the observation that for yeast, where nearly all digenic interactions have been tested, the majority of genes have relatively sparse interaction profiles, while a minority are highly connected (34).

However, we reasoned that appropriate drug perturbations could reveal further dependencies related to the function of the targeted gene. To illustrate this point, we further tested whether compounds that induce DNA damage could reveal additional genetic interactions with the KO of PARP1, a poly-ADP-ribosyltransferase that plays a crucial role in DNA repair mechanisms. PARP1 is highly expressed in NALM6 cells and has been identified as a frequent genetic target in some chemogenomics screens (unpublished data). Because our knockout screens did not reveal any significant PARP1 interactor, we reasoned that induction of DNA damage may increase cell dependency on PARP1 activity, and thus reveal its genetic interactors. To test this hypothesis, we repeated the PARP1 GBGW screen with or without (S)-10-HydroxyCamptothecin (HCPT), a Topo 1 inhibitor (Fig. 4A). Over the eight-day course of drug exposure, HCPT inhibited the growth of both non-targeting controls and PARP1 KO cell populations, but the latter population exhibited greater sensitivity to HCPT (less than 0.5 doublings for both sgRNAs vs 2-3 doublings for non-targeting sgRNAs. Fig. 4D). Chemogenomic screens performed with non-targeting sgRNAs confirmed that PARP1 KO is synthetic lethal (negative scores) to treatment with Topo 1 inhibitors with both sgRNAs (Fig. 4B and unpublished data). Other synthetic lethal hits included the MDR-like drug exporters ABCG2 and ABCC4, both of which are known to contribute to drug resistance by pumping HCPT out of cells (35), as well as components of the homologous DNA repair pathway, while TOP1 was a rescue hit (Fig. 4B). Comparison of the results of the HCPT chemogenomic screen with PARP1 knockout (Fig. 4C, x-axis) vs with control sgRNAs (average of two sgRNAs each) (Fig. 4C, y-axis) indicated that, as anticipated, PARP1 KO was no longer identified as a synthetic lethal hit in PARP1 KO samples in the presence of HCPT (middle of x axis, Fig. 4C), confirming that in these screening conditions it was effectively suppressed. Several other genes identified in the chemogenomic screen with control sgRNAs were also no longer hits when PARP1 was suppressed (upper middle and bottom middle quadrants of Fig. 4D, marked in blue).

**Figure 4:**
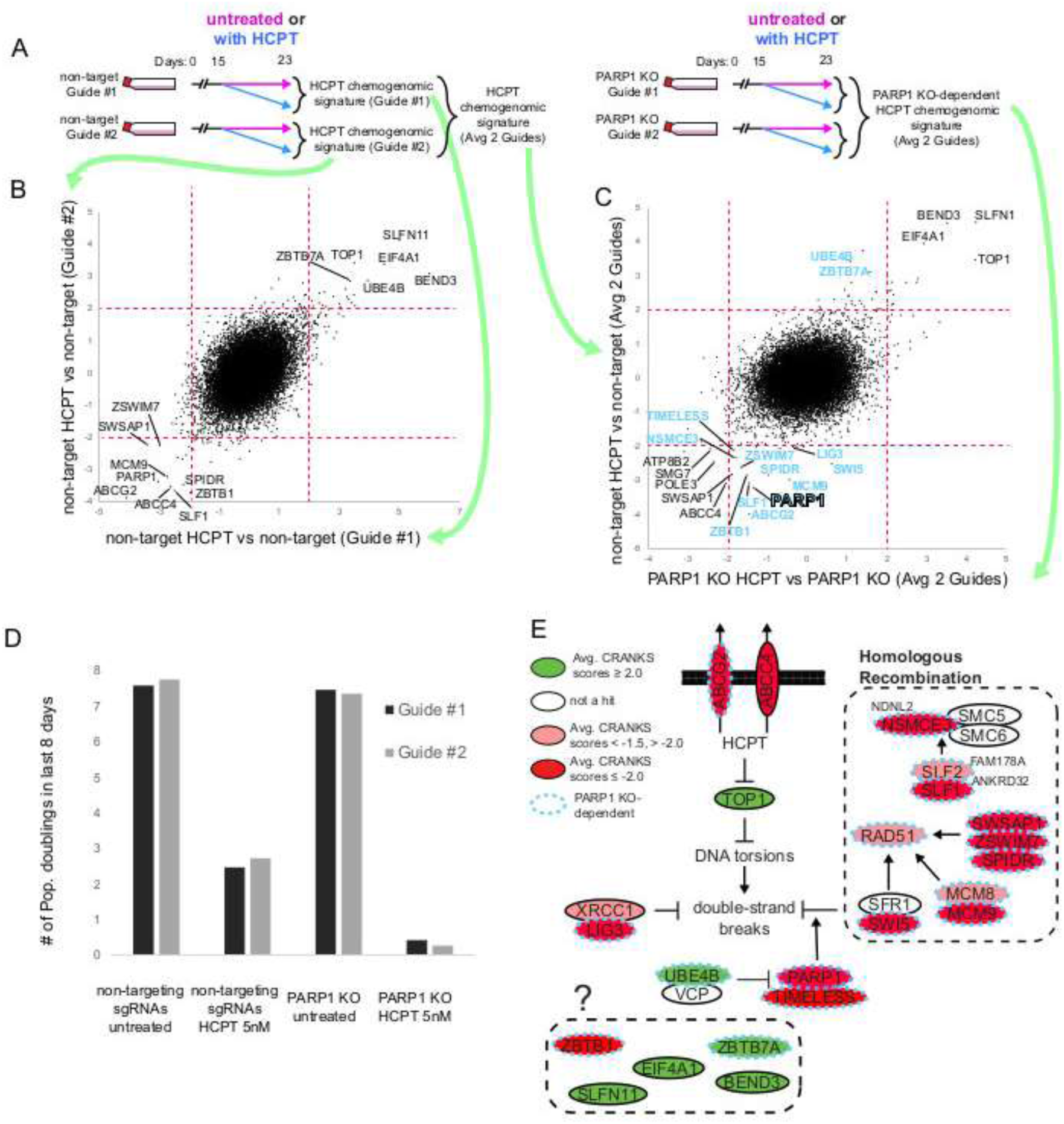
PARP1 genetics revealed by HCTP treatment. **A)** Timeline of GBGW population chemogenomic screens. The timeline is similar to our untreated GBGW screens. However, the final eight days of expansion following Cas9 induction with doxycycline are conducted either with or without chemical treatment. Green arrows indicate where gene scoring events are displayed. **B)** Comparing HCPT chemogenomic screens performed in cell populations infected with each non-target sgRNA provides a robust hit signature. Selected rescue (upper right) or synthetic lethal (lower left) gene KO events are highlighted. **C)** Comparing scores obtained with HCPT chemogenomic KO screens in cell populations infected with PARP1-targeting sgRNA (x-axis, average of both screened guides) with those obtained with similar screens performed in cells infected with non-target sgRNAs (y-axis, average of both guides) shows that hits lost in the in the PARP1 KO population (blue label) include genes mostly involved in HR, in addition to PARP1 itself. **D)** Growth of cell populations during the last eight days of screening with or without HCPT treatment. **E)** HCPT chemogenomic hit genes (red: synthetic lethal, green: rescue) with their current known signalling activities relevant to Topo 1 inactivation by HCPT.

These genes, whose identification as hits is dependent on PARP1 activity (illustrated by dashed blue lines in Fig. 4E), are predominantly linked to homologous recombination, a type of DNA repair in which complexes are recruited to DNA lesions by poly-ADP-ribosylation, which requires PARP1 and TIMELESS (36). On the other hand, other synthetic lethal hits (ABCC4 and other genes in the bottom left corner) or rescue hits (TOP1 and other genes in the upper right corner) were not affected by PARP1 suppression, suggesting that their roles in the response to HCPT is independent of PARP1. Indeed, HCPT poisons TOP1 and is exported by ABCC4 irrespective of the presence of PARP1. These observations indicate that gene vs GW KO screens can be useful comparators to tease out contextual factors to reveal the underlying genetics by activating relevant signaling pathways.

Because **TOP2A** was previously identified as a rescue gene in our chemogenomic screens employing doxorubicin, a Topoisomerase 2 poison (unpublished data), we tested if knocking out TOP2A in the presence of the drug would yield additional mechanistic information on its genetic interactors. TOP2A knockout was protective against doxorubicin-induced growth inhibition, presumably because suppression of the target prevents formation of genotoxic drug-protein complexes (Fig. 5A). The results of the doxorubicin chemogenomic screens conducted with the non-targeting sgRNAs confirmed the previous findings of TOP2A as a robust hit (Fig. 5B). The chemogenomic signature of doxorubicin is enriched in genes implicated in either non-homologous end-joining (NHEJ) or homologous recombination (HR) (Fig. 5C). When analyzing the results of the TOP2A knockout experiments in the presence of doxorubicin (Fig. 5D, x-axis), and comparing them to the chemogenomic signature of doxorubicin (Fig. 5D, y-axis), we observed that TOP2A was lost as a rescue hit in screens with both targeting sgRNAs. Interestingly, in the absence of TOP2A, TOP2B was also lost as a rescue, consistent with Topo 2 heterodimer being important for increasing cell fitness in the presence of doxorubicin (see above and Fig 3). The synthetic lethality of several genes associated with DNA repair was also strongly attenuated, consistent with TOP2A suppression resulting in fewer double stranded breaks. On the other hand, deletion of ZNF451, TDP2 and RAD54L2, involved in resolution of covalent cleavage complexes, remained synthetic lethal in the TOP2A-depleted population, suggesting that these genes, acting independently of TOP2A, lower doxorubicin-induced toxicity by resolving TOP2B- as well as TOP2A-induced covalent cleavage complexes.

**Figure 5:**
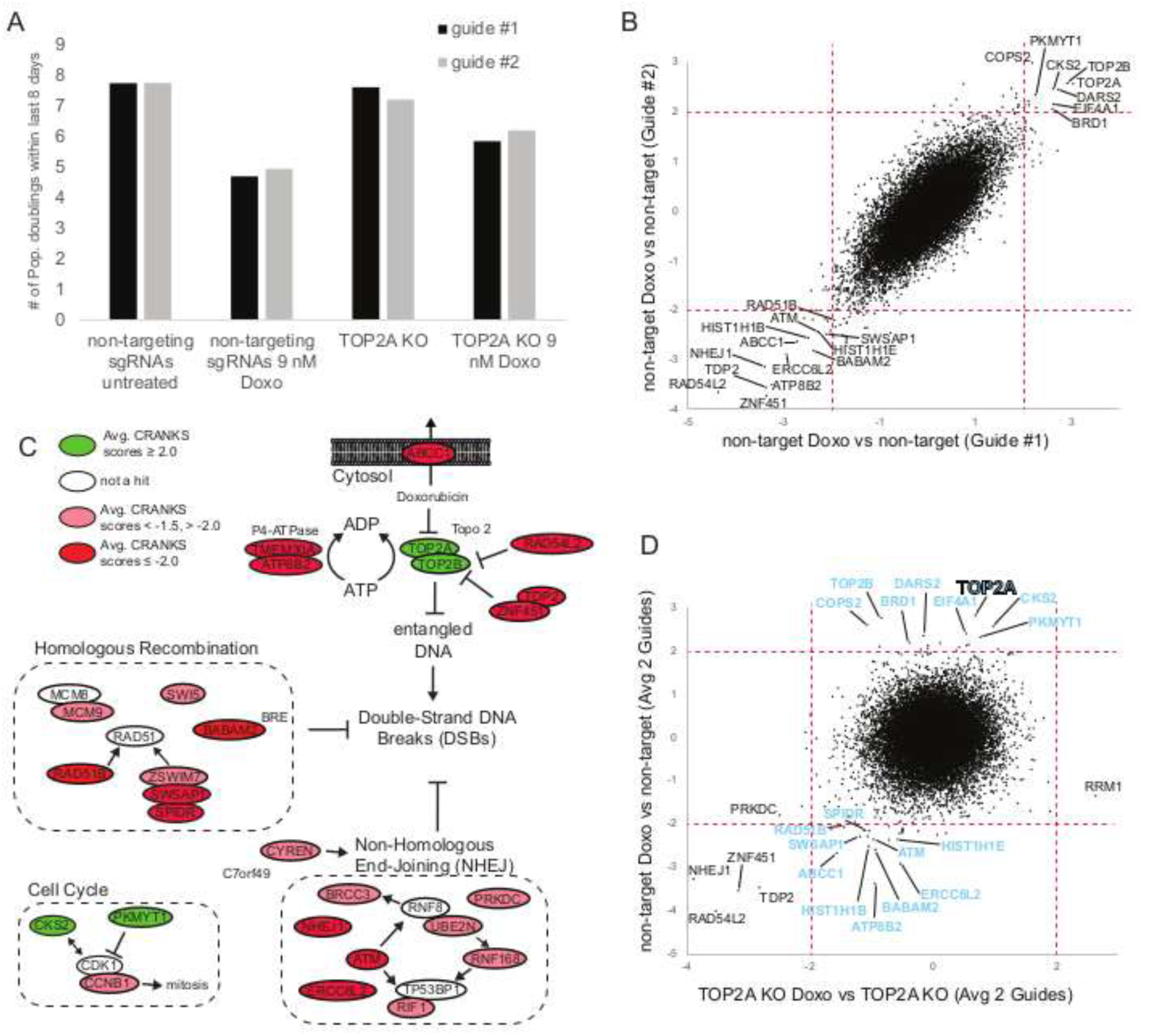
TOP2A genetic interactors revealed by GBGW KO screen with doxorubicin treatment. **A)** Growth of cell populations with or without TOP2A KO during the last 8 days of screening in response to doxorubicin treatment. **B)** Comparing doxorubicin chemogenomic screens performed in cell populations infected with each non-target sgRNA provides a robust hit signature. Selected rescue (upper right) or synthetic lethal (lower left) hits are highlighted. **C)** Doxorubicin chemogenomic hit genes with their current known signalling activities relevant to Topo 2 poisoning by doxorubicin. **D)** The comparison of doxorubicin chemogenomic screens performed in the TOP2A KO cell population arm of a TOP2A GBGW KO screen (x-axis, average of both guides) vs the non-target (y-axis, average of both guides) identifies genes that are no longer hit within the TOP2A KO cell population (blue label), including both TOP2A and TOP2B.

### Gain of Function Screens: MYC

The success of our GBGW loss of function approach based on direct transduction of an existing GW knockout pool by sgRNA expressing lentiviruses described above, encouraged us to assess if a conceptually similar gain-of-function screen could be similarly informative. For this proof-of-concept, we overexpressed MYC, a well-known oncogenic transcription factor, expressed under the control of the strong constitutive promoter MNDU3. Overexpression of mCherry was used as a negative control. We adapted the same experimental framework as for the GBGW screens (Fig. 1A), using MYC-expressing lentiviruses to infect a population of cells already bearing the EKO library. In our experimental setup, MYC overexpression was found to inhibit cell proliferation (Fig. 6A). As evidence that this screen worked as intended, the KO of MYC itself generated the 10^th^ strongest rescue phenotype (Fig. 6B), corroborating that the fitness defect was due to MYC overexpression. The top rescue hit identified was SAE1, a component of an E1 SUMO ligase known to be transcriptionally activated by MYC (37). This finding aligns with earlier studies suggesting that the oncogenic activation of MYC may rely on its ability to enhance SAE1 transcription, thereby creating a potential therapeutic target through SUMO inhibition (38). Collectively, these results indicate that overexpression of MYC could result in increased SUMOylation due to elevated SAE1 levels. Although additional experiments are required to validate these connections, this initial screen illustrates that GBGW GOF screens can be informative and generate new hypotheses.

**Figure 6:**
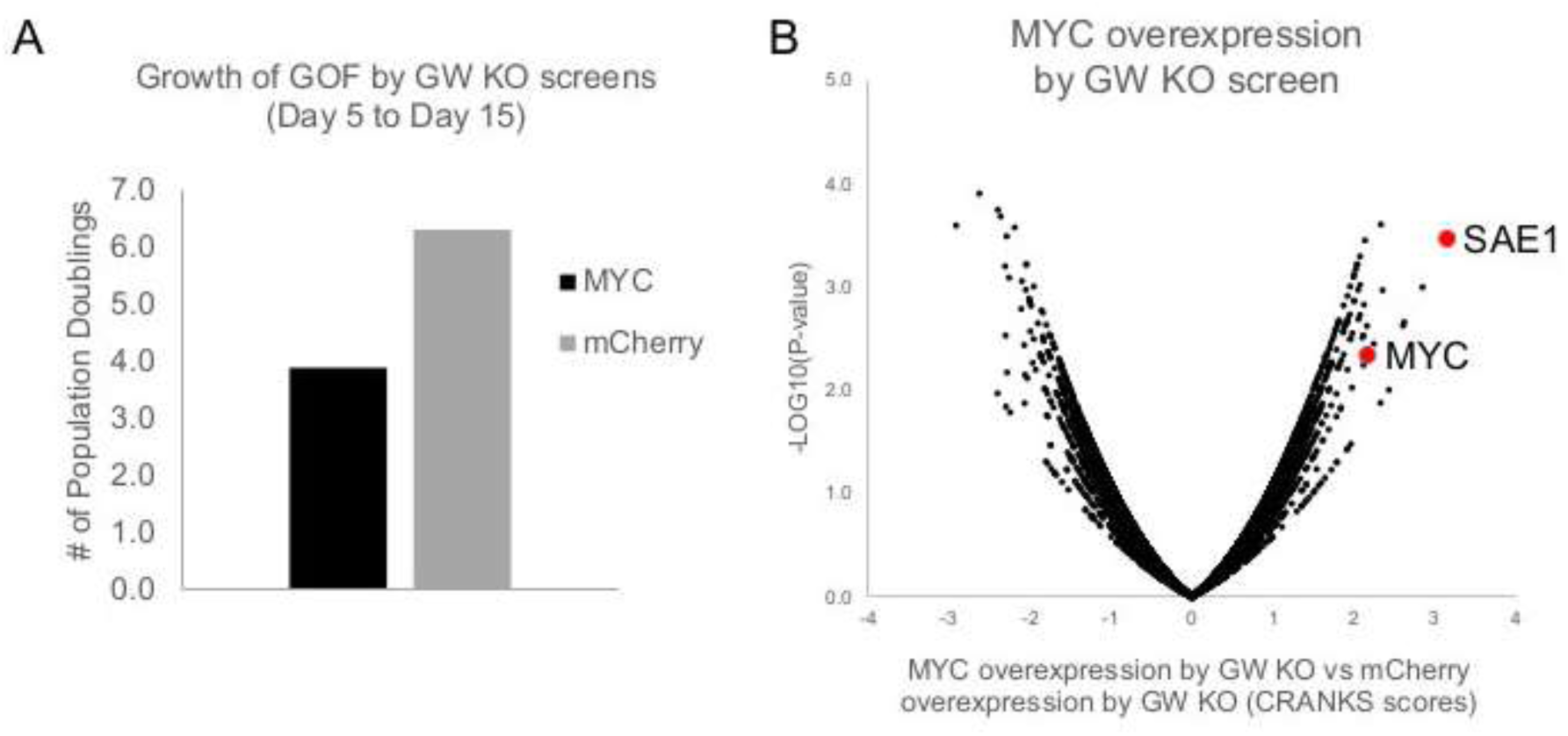
MYC overexpression by Genome-Wide KO screening results. **A)** MYC overexpression-induced toxicity is demonstrated by the marked reduction in population doublings in comparison to mCherry negative control. **B)** Volcano plot of gene scores in the MYC GBGW GOF screen vs mCherry control screen with selected genes highlighted.

## Discussion

The primary objective of this study was to develop a scalable, robust assay capable of systematically evaluating the impacts of double gene knockouts on cellular fitness in human cells. We set out to ensure that the assay was both time and cost efficient; addressing the labor and time constraints associated with the use of custom multiplexed sgRNA libraries or stable knockout cell clones. To achieve this objective, we extended a previously established methodology designed to elucidate the genetic network of UBE2G1 in NALM6 cells (16). In contrast to our previous experimental design, instead of performing the viral pooled GW sgRNA library infection following the transduction of guides targeting the gene of interest, we inverted the sequence of operations by using the uninduced EKO genome-wide knockout library which we routinely employ in chemogenomic screens (39). This approach offers additional advantages over existing methods: it establishes a uniform initial library distribution for all double gene knockout experiments, minimizes the overall number of control samples required for sequencing and facilitates the seamless incorporation of these experiments into the ChemoGenix screening pipeline. Also, this method eliminates the necessity of generating knock-out clones, which can be particularly challenging when dealing with essential gene and bring the risk of secondary mutations that can potentially mask the effects of the original gene knockout.

The results generated in this study indicate that this method effectively identifies established genetic dependencies, even when targeting essential genes such as TOP2A. Other potentially new interactions identified will need further testing to determine their biological significance. Furthermore, also the adaptation of our methodology to GBGW GOF screens can reveal established genetic interactions.

A dearth of identification of genetic interactors may simply reflect the fact that no gene hit combined with suppression of the target affects cell fitness in NALM-6 cells under the conditions of the assay. Nonetheless, it is important to emphasize that the human genome exhibits significant complexity compared to model organisms previously interrogated for genetic interactions, which typically have smaller genomes. Both gene redundancy and transcriptome plasticity further complicate the exhaustive identification of relevant genetic dependencies, while providing cells with the biological buffering necessary to survive diverse stresses.

An analysis of the expression levels of the 8 out of 26 genes that yielded robust hits without chemical treatment revealed that they are significantly more expressed in NALM6 cells than the other 18 genes (Fig. 7A). In contrast, gene essentiality ranking in NALM6 cells does not seem to serve as a reliable predictor for the likelihood of observing genetic interactions for the query gene (Fig. 7B). Detecting genetic interactions for essential genes is expected to be challenging due to the dominant effect of their deletion. Nonetheless, very essential genes such as DDX5, MDM2 and TOP2A produced meaningful genetic signatures. Interestingly, the most effective predictor of success in untreated screens appears to be the frequency with which a gene is identified as a hit in a series of chemogenomic screens we have previously conducted with a diverse array of compounds (Fig. 7C). This high frequency of identification likely reflects several factors, including expression levels and higher genetic connectivity across various chemical perturbations.

**Figure 7:**
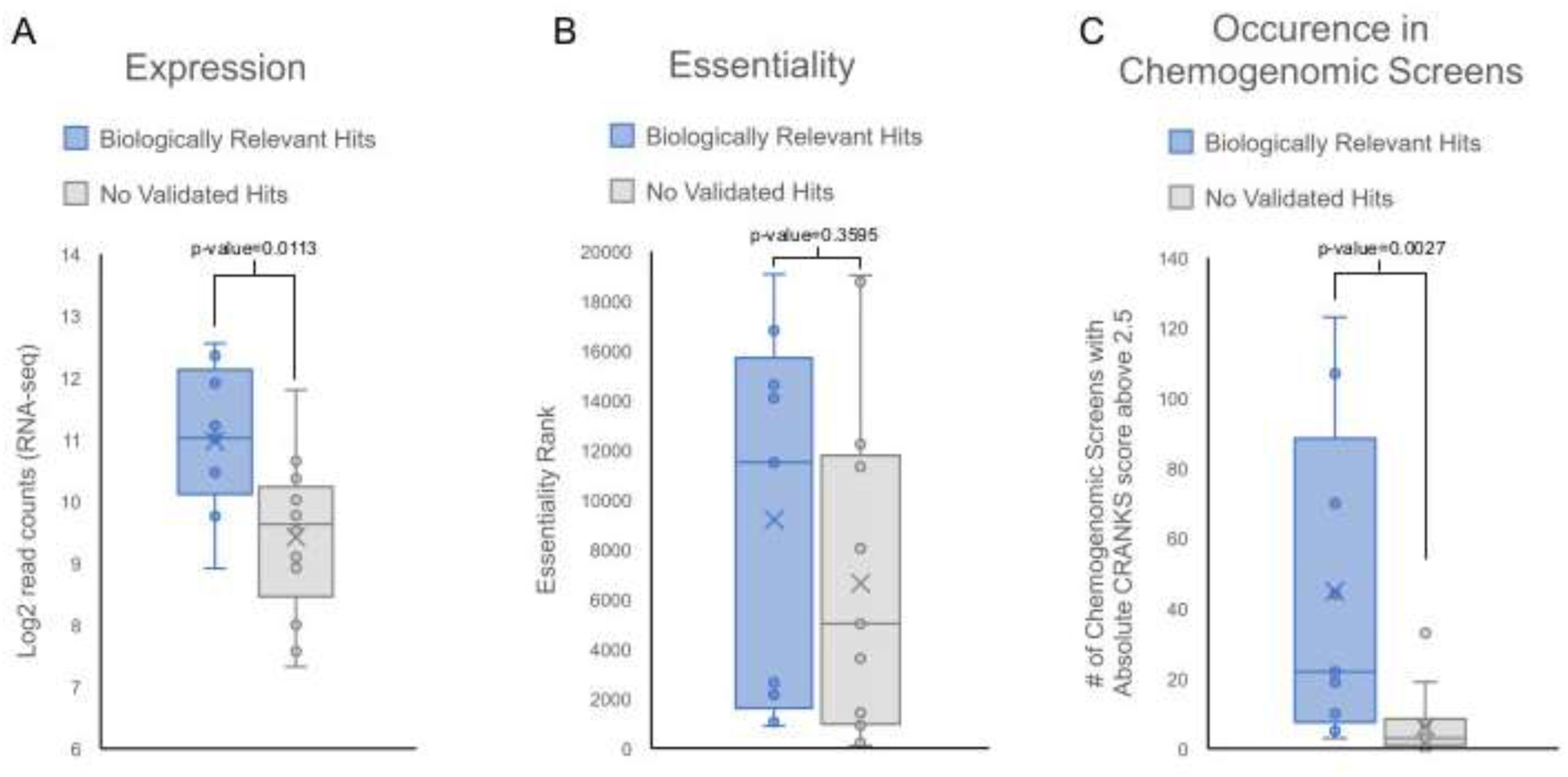
Screen success indicators. Genes targeted in GBGW screens that yielded biologically significant outcomes were assessed according to their expression levels (**A**), gene essentiality (**B**) and the prevalence of these genes as hits with CRANKS scores surpassing absolute values of 2.5 in a wide array of chemogenomic screens (**C**).

Drug treatment can be also exploited to reveal genetic interactors of genes that influence cellular responses to these chemicals. A clear illustration is offered by our studies of the PARP1 gene. Although the initial screening for PARP1 did not yield any significant results, conducting the experiment again in the presence of the TOP 1 inhibitor HCPT revealed its ability to recruit HR components to DNA lesions. This indicates that the specific functions of a gene may need to be perturbed under the screening conditions used for the identification of genetic co-dependencies. GBGW chemogenomic screens can therefore help enrich the landscape of genetic interactors for a given gene of interest.

Although our approach efficiently and rapidly probes all protein-expressing genes for genetic interaction with a gene of interest, we recognize that it may lead to false negatives due to the failure to sample all possible KO combinations with a particular gene of interest. Based on an analysis of all the screens performed to date, we estimate that the gene knockout rate in our dox-inducible NALM-6 clone is close to 80%, but can vary for each sgRNA, typical of CRISPR-Cas9 KO approaches. Double gene knockouts at a population level therefore can be expected to be represented in up to 64% (80% squared) of cells infected by both viruses, reducing the representation of double KOs and increasing noise. The intrinsic variability in the on- and off-target impact of sgRNAs targeting the gene of interest in our GBGW approach can be mitigated by performing paired screens with two or more sgRNAs against the same gene, reinforcing confidence in observed hits. These considerations suggest that our approach is best used early on in a screening campaign, taking advantages of the time and cost efficiencies, after which clonal genotyped approaches can be implemented on a more selective basis for deeper sensitivity.

Another limitation intrinsic to all screening experiments is the cell model used for these experiments. NALM6 cells are leukemic cells that carry an array of genetic defects, often of little known biological importance. Although all screens were carried out in this model, for which we already had constructed a genome-wide knockout pool library, the approach described herein may be used in other cell lines more suitable for specific biological questions.

## Material & Methods

### Cell lines

The NALM6 pre-B lymphocytic human cell line was kindly supplied by Prof. Steve Elledge (Harvard University). NALM6 pCW-Cas9 clone #20 was generated using lentivirus derived from the plasmid pCW-Cas9 (Addgene # 50661), followed by puromycin selection and FACS sorting as described (3). Clone #20 underwent further infection with lentiviruses expressing our EKO sgRNA pooled library and selection with blasticin. HEK293T cells were sourced from ATCC (CRL-3216). Cells were cultured in RPMI (NALM6) or DMEM (HEK293T) supplemented with 10% FBS (Sigma) at 37°C in a 5% CO2 incubator.

### Plasmid cloning

The sgRNA sequences were selected using the CRISPick tool (https://portals.broadinstitute.org/gppx/crispick/public). The two sgRNA sequences that received the highest scores were chosen for further application. sgRNA sequences were cloned into the pLX-sgRNA-2X-BsiWI-AAVS1-Hygro plasmid developed internally through Gibson assembly by modifying pLX-sgRNA (Addgene #50662). The AAVS1 sequence was removed by digestion with BsiWI followed by gel purification. A 60-61 bp oligonucleotide was used as a template for PCR with oligos FWD GA sgRNA -50 to 0 and REV GA sgRNA 0 to +50. The resulting PCR product was then cloned by Gibson Assembly into the BsiWI-digested plasmid.

mCherry was amplified through PCR from a plasmid provided by Étienne Gagnon (IRIC, Université de Montréal) using the primers “FWD mCherry-3XFLAG for GA MNDU3” and “REV mCherry-STOP for GA MNDU3.” MYC cDNA was amplified from a plasmid provided by Philippe Roux (IRIC, Université de Montréal) using the primers “FWD cMYC for GA MNDU3” and “REV cMYC for GA MNDU3”. Both mCherry and MYC were amplified employing KAPA HiFi HotStart DNA Polymerase (Roche) and cloned by Gibson Assembly into the EcoRI-digested pCCL-c-MNDU3-PGK-Hygro plasmid derived from the lentiviral vector pCCL-c-MNDU3-X (Addgene #81071) to enable hygromycin selection. Following transformation into STBL3 *E. coli*, midipreps were performed using the PureLink kit (Invitrogen), and guide sequence insertions were confirmed through Sanger sequencing.

### Lentivirus production

Lentiviruses were produced by co-transfecting two helper plasmids, psPAX2 (Addgene #12260) and pCMV-VSV-G (Addgene #8454), along with plasmids encoding sgRNAs in HEK293T cells, as previously described (3). psPAX2 (6 μg), pCMV-VSV-G (3 μg) and pLXsgRNA-Hygro (9 μg each) were diluted step-wise in 667 uL water supplemented, after vortexing, with 53 ul of polyethyleneimine 1 mg per mL. The resulting mixture was added dropwise over 80-90% confluent HEK293T cells in 10 mL fresh media in a 10 cm dish. After 16 hours, the media was replaced with 2% FBS DMEM, and 48 hours later, the lentivirus-containing supernatant was collected, filtered through a 0.45 μm filter, and combined with a filter-sterilized (0.22 μm) concentrated stock of preservation solution to achieve a final concentration of 5% sucrose, 2 mM MgCl2, and 10 mM HEPES at pH 7.2. The harvested lentiviral preparations were subsequently stored at -80°C until they were used for infecting the EKO library introduced in NALM6 pCW-Cas9 clone #20.

### Gene by GW double CRISPR knockout screens

The screens were conducted in three distinct rounds. Each round used 8 x 5mL cryovials containing 22.5E6 NALM6 pCW-Cas9 clone #20 (total of 180E6 cells) transduced with the EKO sgRNA library (“first sgRNA”) and frozen in 4.5 mL 50% FBS, 40% RPMI and 10% DMSO. Following a centrifugation step at 330g for 7 minutes and resuspension in media, the cells were quantified using a Z2 particle count and size analyzer (Coulter counter; Beckman) and resuspended in media at 400,000 cells per mL. The cells were then expanded in a spinner flask (Corning) under continuous slow agitation (20 turns per minute) for 5 days, undergoing exponential growth dilution and ultimately achieving a target concentration of 2 million cells per mL on the final day. A total of 140 mL of cells, equating to 280 million cells, was centrifuged for 7 minutes at 330g, and the resultant cell pellet was processed using the Puregene (Qiagen) genomic DNA extraction kit to create a Day 0 sample control. The remaining cells in the spinner flask were treated with 1000X filter-sterilized 10 mg/mL Protamine Sulfate and incubated with gentle swirling for 20 minutes. Subsequently, T-175 flasks were inoculated with either 140 mL of cells (for GBGW screens with a representation of 250X cells per “second” sgRNA; e.g. PTEN, TP53, MDM2, NPRL3, ADK, HMGCL, or PARP1 screens) or 70 mL of cells (for GBGW screens with a representation of 100X cells per sgRNA; e.g. PTEN, PARP1, BANF1, CCNQ (FAM58A), CDADC1, CSE1L, DDX3X, DDX5, ERCC2, ERCC3, ETFDH, FAM172A (ARB2A), LONP1, MAPK14, PLCXD1, RBM26, SMARCD1, TOP2A, TOP2B, YME1L1, or ZNF451 screens). These were either left untreated (no second sgRNA control) or supplemented with 30 mL (250X representation) or 15 mL (100X representation) of second sgRNA-carrying lentivirus preparation. Under these conditions, the MOI was estimated to be close to 0.5 as assessed by parallel infection with a lentivirus constructed with LentiCRISPRv2GFP (Addgene #82416) and subsequently measuring two days later the percentage of GFP+ cells by FACS. On the first day, the 250X screens were divided between two T-175 flasks, with the total volume increased to 340 mL (170 mL in each flask) by adding media. Each time the 250X pooled screens required dilution and expansion to a volume exceeding 170 mL, the contents of both T-175 flasks were combined, thoroughly mixed, diluted, and subsequently redistributed into two separate T-175 flasks. On Day 1, the 100X screens received 85 mL of fresh media, resulting in a total volume of 170 mL within a single flask. On Day 2, half of the cells from each pooled screen were discarded, and the volumes were adjusted back to the original volume by adding media containing hygromycin to achieve a final concentration of 300 µg/mL. On Day 5, cell concentrations were measured using a Z2 Coulter counter, and 272 million (250X) or 136 million (100X) cells were re-seeded at a density of 800,000 cells per mL, resulting in final volumes of 340 mL and 170 mL, respectively, while still under hygromycin selection. On Days 8 and 11, cell concentrations were again assessed, and 102 million (250X) or 51 million (100X) cells were re-seeded at 600,000 cells per mL, with total volumes of 170 mL and 85 mL, respectively. On Day 8, hygromycin was included, whereas on Day 11 it was omitted, but the media was supplemented with 2 µg/mL (final concentration) of doxycycline hyclate (Sigma). On Days 13 and 15, cell concentrations were measured once more, and 68 million (250X) or 28 million (100X) cells were re-seeded at a density of 400,000 cells per mL, with total volumes of 170 mL and 70 mL, respectively, with doxycycline on Day 13 and without it on Day 15. For screens requiring chemical treatments beginning on Day 15, the number of cells seeded on Day 13 was increased 2 folds to ensure an adequate supply of cells by Day 15. Compounds were prepared in DMSO at a concentration 1000 times higher than the dose anticipated to achieve 50% growth inhibition, with 1/1000 of this volume (0.1% v/v DMSO) administered to the treated samples. On Days 17, 19, and 21, cell counts were conducted, and if the concentration exceeded 800,000 cells per mL, the cells were diluted back to 400,000 cells per mL (with a total volume of 170 mL for 250X or 70 mL for 100X). Additional chemicals were added based on the volume of media added, assuming that the compounds within the cells remained active. If the cell concentration was below 800,000 cells per mL, no dilution was performed. On Day 23, cell counts were taken, and 100 million cells (for 250X) or 50 million cells (for 100X) were pelleted at 330 g for 7 minutes, followed by genomic DNA extraction utilizing the Puregene kit.

### Next generation sequencing

sgRNA sequences were amplified in an initial round of PCR using 462 µg of genomic DNA (250X representation, equivalent to 70 million cells). The reaction mixture consisted of 575 µL of 10X PCR buffer, 115 µL of 10 mM dNTPs, 23 µL of 100 µM Outer primer 1, 23 µL of 100 µM primer BLAST FWD1, 115 µL of DMSO, and 145 units of EasyTaq DNA polymerase (Transgene Biotech), resulting in a total volume of 5.75 mL. Alternatively, 185 µg of genomic DNA (100X representation, or 28 million cells) was used as a template, employing the same PCR recipe but in volumes reduced by a factor of 2.5. The BLAST FWD1 oligonucleotide is specifically designed to hybridize within the blasticidin selection marker, which is exclusive to the EKO GW sgRNA library, thereby preventing amplification and sequencing of the second sgRNA intended for double gene knockout which utilizes a hygromycin selection marker. In instances where the required DNA quantities were not achieved following genomic DNA extraction, all available genomic DNA was used as template for the PCR, and an appropriate number of additional PCR cycles were conducted. Multiple 100 µL reactions were prepared in a 96-well format on a T100 thermal cycler (BioRad, Hercules, California, USA) with the following cycling conditions: 95°C 5 min, 26 cycles of 35 sec at 94°C, 50 sec at 52°C and 1 min at 72°C, final step of 10 min at 72°C after the last cycle. The completed reaction mixtures were pooled into a single tube and mixed thoroughly. Following confirmation of amplification of a 718bp fragment containing the sgRNA sequence on a 1% 1X TAE agarose gel, a second PCR reaction was performed to add Illumina sequencing adapters and 6 bp indexing primers. 10 ul of 1:20 dilution of unpurified PCR1 product were PCR-amplified using: 10 µl 5X Kapa buffer, 5 µl 2,5 mM dNTPs, 1 µl of 100 µM TruSeq Universal Adapter +0 (IDT Ultramer), 1 µl of 100 µM TruSeq Adapter with appropriate index (IDT Ultramer), 1 µl DMSO and 5 units Kapa HiFi HotStart DNA polymerase and volumes brought to 50 µl total volume (5 min at 95°C, 5 cycles of 15 sec at 95°C, 30 sec at 50°C and 30 sec at 72°C, 5 cycles of 15 sec at 95°C, 30 sec at 56°C and 30 sec at 72°C, followed by 5 min final step at 72°C after the last cycle). After verifying the generation of a 238 bp amplicon on a 2% 1X TAE agarose gel, with each band exhibiting equal abundance, 5 µl or 10 µl from each sample intended for sequencing on the same lane were pooled and purified using solid-phase reversible immobilization (SPRI) beads (AxyPrep Fragment Select-I Kit) at a 2:1 ratio of SPRI beads to PCR product. The samples were sequenced on an Illumina NextSeq 500 with a 75 cycle single-read high output flow cell, capturing a total of 43 bp, with the first 23 bp reads occurring in dark cycles (IRIC’s Genomic core facility, Montreal, Quebec, Canada).

### Gene Enrichment/Depletion Scoring

Gene interaction scores were obtained using a modified version of the RANKS (3) algorithm, which utilizes guides targeting genes of similar essentiality as controls. This methodology allows for the distinction between condition-specific chemogenomic interactions and non-specific fitness or essentiality phenotypes. As background for the analysis, we summed the reads obtained from the two samples expressing independent control sgRNAs targeting the AAVS1 locus.

## Data availability

All the data presented in this work is freely available on the ChemoGenix website (URL: https://chemogenix.iric.ca) where forthcoming experiments will also be made available.

## Acknowledgements

We acknowledge the Fonds de recherché du Québec (FRQ) – secteur Santé for its support of the Institute for Research in Immunology and Cancer (IRIC), an FRQ-designated center. This work was supported by grants from the Canadian Institutes of Health Research [grant PJT-173443 to J.A.L., grant PJT-191684 to N.P., grant PJT-190250 to S.M., grant FDN-167277 to M.T., Tier 1 Canada Research Chair in Translational Genomics to C.N., Tier 1 Canada Research Chair in Regulation of mRNA Translation and Metabolism to I.T.], The Cancer Research Society [grant #1281387 to J.D.N.], NSERC Discovery [grant RGPIN-2022-04206 to V.A., grant RGPIN-2020-04947 to B.T.W.].

---

**Supplementary Figure 1.**
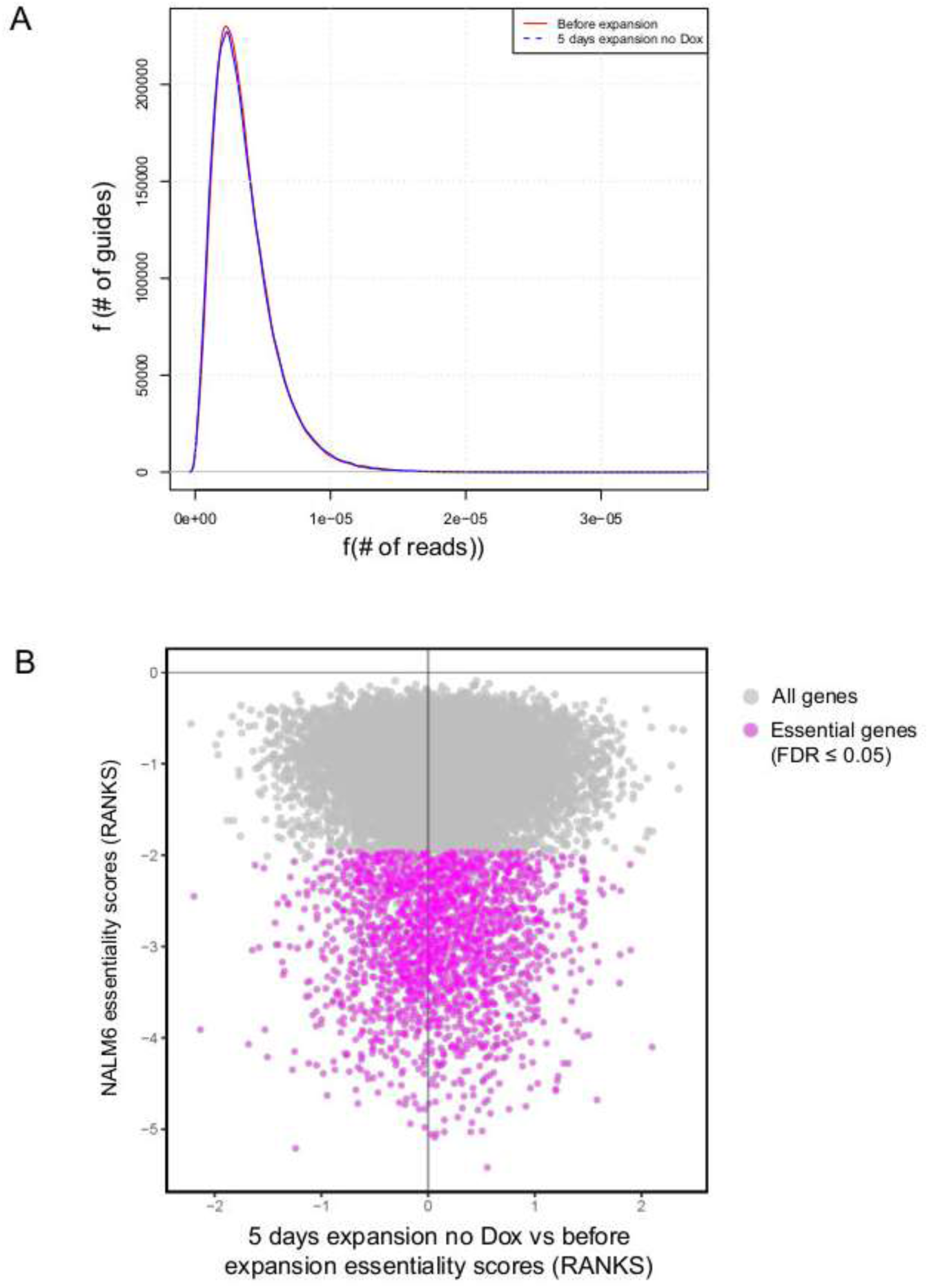
A) The representation of sgRNAs remains largely consistent from day -5 (red curve) to day 0 (purple curve). **B)** A five-day pre-expansion of the pooled GW sgRNA library in a spinner flask without doxycycline treatment does not result in the loss of sgRNAs targeting essential genes from the pool.

**Supplementary Figure 2.**
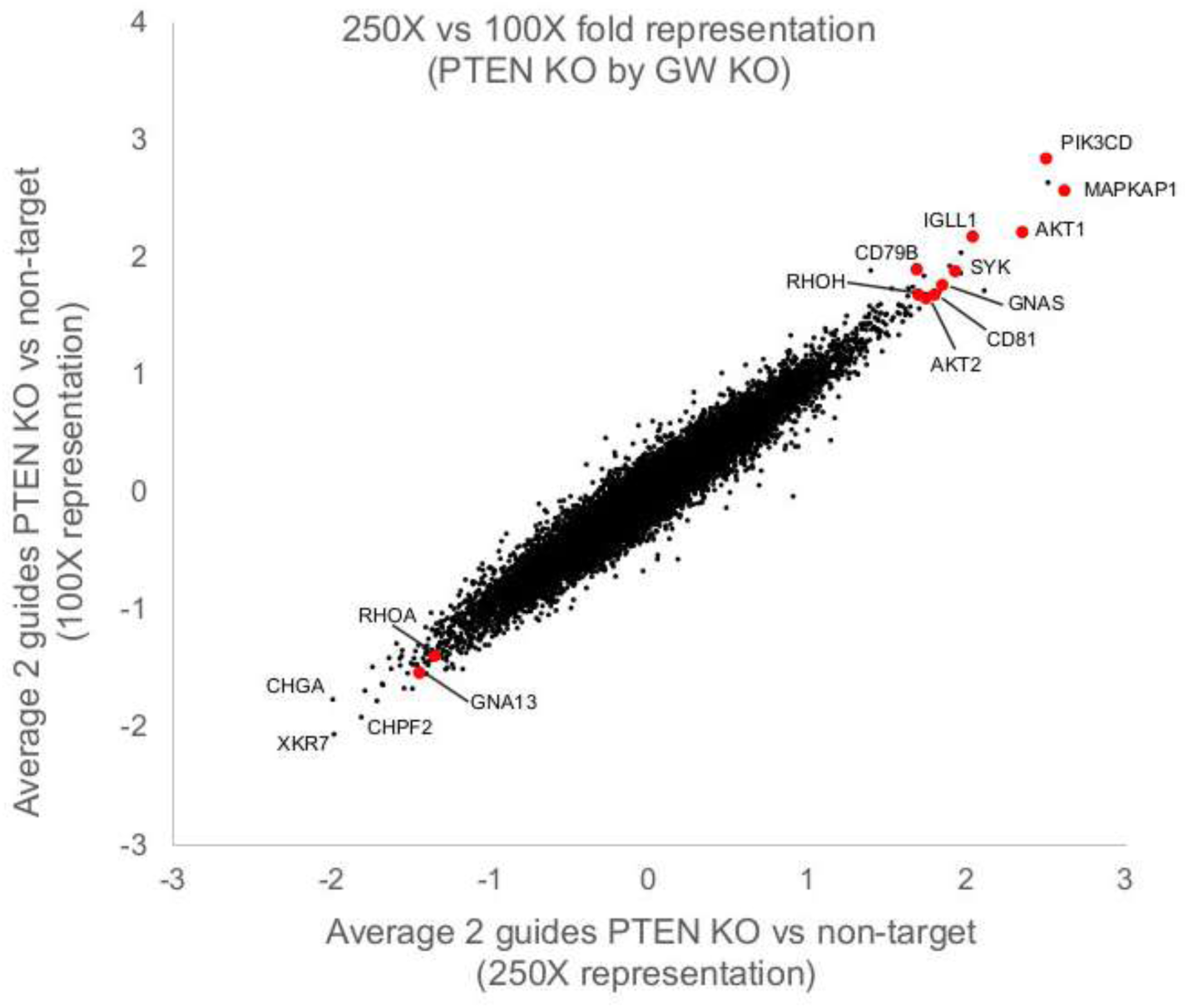
PTEN GBGW KO screens screened at 250X or 100X fold representation (# of cells per sgRNA) generate largely the same genetic signature.

## Notes

### Competing Interest Statement

The authors have declared no competing interest.

